# Red queen’s race: rapid evolutionary dynamics of an expanding family of meiotic drive factors and their hpRNA suppressors

**DOI:** 10.1101/2021.08.05.454923

**Authors:** Jeffrey Vedanayagam, Ching-Jung Lin, Eric C. Lai

## Abstract

Meiotic drivers are a class of selfish genetic elements that are widespread across eukaryotes. As their activities are often detrimental to organismal fitness, opposing regulatory mechanisms are usually required to silence them, to ensure fair segregation during meiosis. Accordingly, the existence of such selfish elements is frequently hidden in genomes, and their molecular functions are little known. Here, we trace evolutionary steps that generated the *Dox* meiotic drive system in *Drosophila simulans* (*Dsim*), which distorts male:female balance (*sex-ratio*) by depleting male progeny. We show that *Dox* emerged via stepwise mobilization and acquisition of portions of multiple *D. melanogaster* genes, notably including from protamine, which replaces histones in haploid sperm and mediates the highly condensed state of sperm chromatin. Moreover, we reveal novel *Dox* homologs in *Dsim* and massive, recent, amplification of *Dox* superfamily genes specifically on X chromosomes of its closest sister species *D. mauritiana* (*Dmau*) and *D. sechellia* (*Dsech*). The emergence of Dox superfamily genes is tightly associated with 359-bp repeats (in the 1.688 family of satellite repeats) that flank *de novo* genomic copies. In concert, we find coordinated emergence and diversification of autosomal hairpin RNA-class siRNA loci that target subsets of *Dox* superfamily genes across *simulans* clade species. Finally, an independent set of protamine amplifications on the Y chromosome of *D. melanogaster* indicates that protamine genes are frequent and recurrent players in sex chromosome dynamics. Overall, we reveal fierce genetic arms races between meiotic drive factors and siRNA suppressors associated with recent speciation.

## Introduction

Meiotic drive is widespread in plants, animals, and fungi (Agren and Clark, 2018; Zanders and Unckless, 2019). Yet, we know very little about their origins, molecular functions, and short and long-term persistence in wild populations (Lindholm et al., 2016). A particular type of meiotic drive is sex chromosome drive, which occurs when transmission of sex chromosomes (XY or ZW) deviates from expected Mendelian segregation. This frequently manifests in the heterogametic sex, resulting in unequal transmission of the X and Y chromosomes (or alternatively, the Z and W chromosomes) during gamete formation (Jaenike, 2001). In sexually reproducing species, the male-to-female ratio amongst progeny is roughly equal (Fisher, 1930). However, during *sex-ratio* (SR) drive, the unequal transmission of sex chromosomes typically impairs the representation of sons. SR drive systems occur broadly across eukaryotic species and are widely reported in multiple *Drosophila* species, indicating their recurrent and facile emergence (Jaenike, 2001). While there is to date little evidence to indicate any strong SR drive loci in the model fruitfly *D. melanogaster* (*Dmel*), genetic studies in its sister species *Drosophila simulans* (*Dsim*) revealed at least three different SR drive systems. These were named based on their location of geographical isolation and/or primary study, and are known as the Paris (Helleu et al., 2016), Durham (Tao et al., 2001) and Winters (Tao et al., 2007a; Tao et al., 2007b) SR systems. Together, these studies highlight that alleles involved in SR drive and suppression can evolve with extraordinary dynamics, and that multiple SR systems can co-exist in an individual species simultaneously.

*Dsim* and its immediate sister species *D. sechellia* (*Dsech*) and *D. mauritiana* (*Dmau*) comprise the *simulans* clade, which diverged from a *Dmel*-like ancestor only ~250,000 years ago (Garrigan et al., 2012). As hybrid crosses between *simulans* clade species yield fertile females, albeit sterile males, these species comprise prime models to discover speciation factors using introgression genetics (Masly and Presgraves, 2007; Presgraves et al., 2009). In addition to identifying SR drive systems, such approaches also uncovered hybrid sterility factors in *Dsim*, all of which preferentially or specifically disrupt spermatogenesis (Meiklejohn and Tao, 2010). The Durham drive system was uncovered during introgressions between *Dsim* and *Dsech*, where an ~80Kb minimal autosomal genomic region was inferred to harbor a dominant suppressor of SR drive (*Too much yin*, *Tmy*). In turn, *Tmy* was hypothesized to silence a still-unknown driver gene, whose activity in biasing *sex-ratio* and/or inducing sterility, is cryptic in contemporary *Dsim* populations due to its strong suppression (Tao *et al*., 2001). Subsequently, the Winters SR system was defined by a distinct suppressor locus termed *Not much yang* (*Nmy*) (Tao *et al*., 2007b). The target of Nmy was identified by genetic criteria as the X-linked locus *Distorter on the X* (*Dox*). Mutation of *Dox* bypasses the need for wild-type *Nmy* alleles, since *dox; nmy* double mutants restore equal *sex-ratio* and exhibit normal spermatogenesis (Tao *et al*., 2007a).

Complex evolutionary relationships were already apparent from these early molecular studies (Tao *et al*., 2007a; Tao *et al*., 2007b), since *Nmy* encodes an apparent retroposed copy of *Dox* that forms an inverted repeat. In addition, *Dox* was found to have a paralog termed *Mother of Dox* (*MDox*). All of these *Dsim* loci are absent from the *Dmel* genome, suggesting their likely emergence in *Dsim* or the *simulans* clade ancestor. However, there have not been subsequent insights into the potential molecular activities or evolutionary origins of these meiotic drive genes. In part, such efforts were hindered by subpar genome assemblies in *Dsim*. Sequences with homology to *Dox/MDox* were found on unassembled contigs in the original public genomes, but their chromosomal locations remained ambiguous (Lin et al., 2018). Nevertheless, subsequent *de novo* genome assemblies from short-read data of *simulans* clade species helped infer recent introgression of *Dox/MDox* genes between *Dmau* and *Dsim*, indicating their presence outside *Dsim* (Meiklejohn et al., 2018).

Recently, PacBio genome assemblies from the *simulans* clade were reported, providing an improved resource for evolutionary studies of SR systems across the *simulans* clade (Chakraborty et al., 2021). We previously used the unassembled long-read sequencing data to demonstrate that *Nmy* generates hairpin RNA (hpRNA)-class siRNAs, which are generated by the endogenous RNAi pathway (Czech et al., 2008; Kawamura et al., 2008; Okamura et al., 2008; Wen et al., 2015a). Because Nmy and Tmy hpRNAs comprise nearly perfect inverted repeat structures, they are misassembled in multiple short read *Dsim* assemblies, but are present within continuous PacBio *Dsim* long read sequences (Lin *et al*., 2018). We further showed that *Nmy* silences the *Dox/MDox* driver genes *in trans* (Lin *et al*., 2018). Furthermore, our analyses of small RNAs identified another long inverted repeat within the ~80Kb genomic window bearing the genetic suppressor (*Tmy*) for the Durham SR drive system (Lin *et al*., 2018; Tao *et al*., 2001). Remarkably, the Tmy and Nmy hpRNAs bear substantial homology, suggesting that even though the Winters and Durham systems are genetically separable, they are evolutionarily linked. Furthermore, they hint at the existence of as-yet unidentified meiotic drive genes that are controlled by these hpRNAs, in particular by Tmy.

To recap, prior analyses suggest that *Dsim* Dox distorts normal sex-ratio by depleting Y-bearing sperm, and that this is countered by dominant silencing mediated by complementary siRNAs derived from the hpRNA Nmy. Furthermore, the homology of Nmy with the Tmy hpRNA implied an unappreciated co-evolutionary arms race with an unknown drive factor(s). Since none of these driver or hpRNA suppressors exist in *Dmel*, we sought to distinguish whether Dox emerged de novo, or could be assigned to shared genic origins in *Dmel*. In addition, we sought to delineate the evolution of the Dox system in other *simulans* clade species, since prior introgression studies implied that they were specific to *Dsim*. Taking advantage of the PacBio assemblies, we perform detailed evolutionary analyses of the evolution of Winters and Durham SR systems. We are able to (1) trace the origin of *Dox* from its constituent genes in *Dmel*, including from protamine, (2) uncover rampant proliferation of *Dox* superfamily loci on X chromosomes across the three *simulans* clade species, (3) identify a likely strategy by which *Dox* superfamily loci expand and mobilize via flanking satellite repeats, and (4) show that the dynamics of *Dox* superfamily meiotic drive loci go hand-in-hand with the diversification of hpRNA loci that bear homology to various subsets of the *Dox* superfamily. These findings clarify the evolutionary links between Winters and Durham SR systems, and testify to ongoing genetic arms races in the *simulans* clade. Moreover, the rapid evolutionary dynamics of SR drive elements and their suppression by RNAi loci highlight the role of endo-siRNAs in resolving intragenomic conflicts and potentially acting as agents of speciation.

## Results

### The *de novo Dsim Dox* locus is a chimera of multiple genes, including protamine

Yun Tao reported that the *Dsim* meiotic drive locus *Dox* results from an insertion into a genomic region syntenic with *Dmel*, and that *Dox* is homologous to another X-linked locus termed *Mother of Dox* (*MDox*) (Tao *et al*., 2007a). However, as *Dox*/*MDox* seemed to contain only short open reading frames lacking apparent known domains, it was unclear if *Dox* was coding or non-coding (Tao *et al*., 2007a). Moreover, it was not addressed if *Dox*/*MDox* originated fully *de novo*, or alternatively, emerged from existing genes.

With our recent interest in Dox function and its suppression by hairpin RNA (hpRNA) substrates of the endogenous RNAi pathway (Lin *et al*., 2018; Wen *et al*., 2015a), we re-assessed potential functional products of *Dox*. As defined by RNA-seq data from testis *Dsim* (Lin *et al*., 2018), the *Dox* locus encodes a 4.1kb spliced transcript (**Figure 1A**). We systematically searched for similarities at the nucleotide or coding level in *Dsim* and the *Dmel* sister genome, using relaxed parameters to capture possible relationships. Surprisingly, this revealed complex, chimeric origins of *Dox*, fusing sequences clearly identifiable from multiple protein-coding genes and repetitive DNA. In the following section, we list homologies to *Dmel* loci (*CG####* or common gene names) and/or *Dsim* loci (*GD####*). However, to be clear, for nearly all these loci, *Dmel* exhibits the ancestral state and lacks the sequence insertions that are present in several *Dsim* homologs.

**Figure 1.**
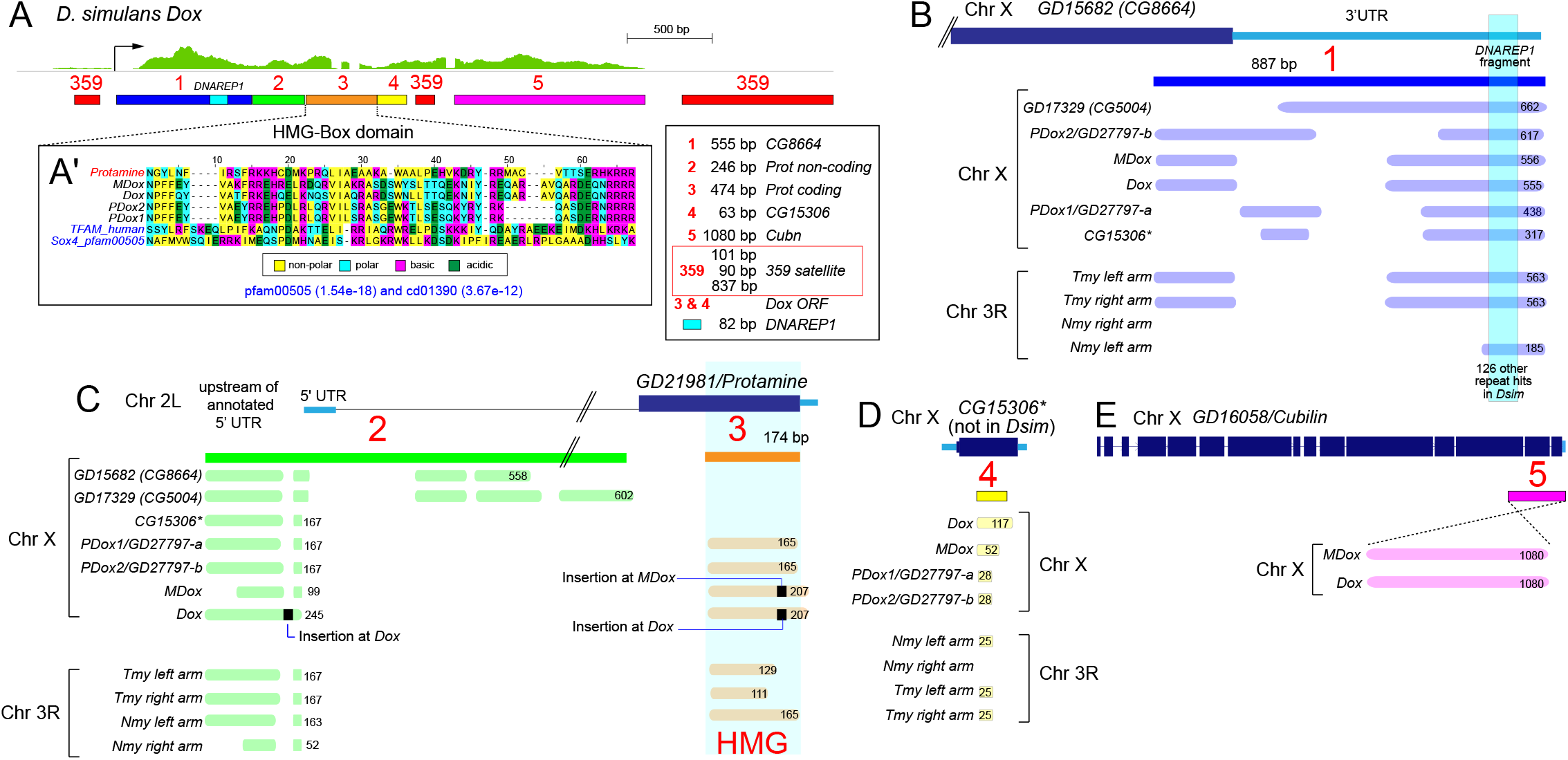
Structure of *Dox* transcript with segments acquired from various genes on the path to its origin. (**A**) Testis RNA-seq data shows a multi-exonic transcript from the *Dox* region, which several distinct segments acquired from protein-coding genes and repetitive elements. 359 corresponds to sequence with similarity to the 359 (also known as 1.688 family) satellite repeat. Segment 1 (blue) corresponds to sequence acquired from *GD15682* (*CG8664*); embedded within this segment is 82 bp derived from DNAREP1 transposable element (turquoise). egments 2 (green) and 3 (orange) correspond to sequences acquired from *GD21981* (*Protamine*). Segment 3 is from the protein-coding portion of Protamine, which harbors an HMG-box domain. Inset (**A1’**) highlight amino acid identify/similarities between Dox family genes and both the Protamine HMG-box domain, and more distant HMG-box sequences from human (pfam definition) and Sox4 (pfam00505) as an outgroup. Segment 4 (yellow) was acquired from *CG15306*, and segment 5 (pink) derives from *Cubilin*. The key depicts Dox segment features, including their segment number, nucleotide length, and origin. (**B**) Overlap of various genomic regions to *GD15682* (*CG8664*) from BLAST search. Segment 1 (blue) corresponds to 887bp from C-terminus and 3’ UTR of *GD15682* (*CG8664*). BLAST hits of various lengths to different genomic features on Chr X and Chr 3R are shown as light blue bars with lengths of nucleotide homology indicated. 82 bp of Segment 1, which corresponds to *DNAREP1* transposable element, retrieves 126 BLAST hits in the *Dsim* PacBio genome (**C**) Genomic matches to the ancestral *protamine* (*GD21981*) gene, include regions with similarity to its upstream/5’ UTR/intronic regions (green), and others bearing the HMG box domain (orange). (**D**) Segment 4 from Dox was acquired from *CG15306*. *CG15306* is no longer extant in *Dsim*, but relics from the insertion can be identified from BLAST search at Dox superfamily genes and their hpRNA suppressors. (**E**) Segment 5 from *Dox* was acquired from C terminus of *Cubilin* (pink). This segment is found only at *Dox* and *MDox*. Note that *Cubilin* matches to the antisense strands of *Dox* and *MDox*.

We divide the *Dox* genomic region based on similarities to known genomic features (**Figure 1A**). We note that the transcription unit is flanked by sequences corresponding to 359 satellite repeats (101 bp upstream and 837 bp downstream), with an additional 90 bp upstream and downstream of 359 repeat found within the transcribed region. 359 repeats belong to the complex 1.688 satellite repeat family in *Drosophila*. In *D. melanogaster*, a large block of 359 satellite repeat is found in the pericentromeric heterochromatin on the X chromosome (Usakin et al., 2007). Whole-genome sequencing from the *simulans* clade revealed recent expansion of 359 satellite repeats on the X chromosome, including a large block within euchromatin (Garrigan et al., 2014).

The 5’ end of the transcription unit includes 555 nts with similarity to the C-terminal-encoding and 3’ UTR regions of *CG8664/GD15682* (designated “1”). Embedded within this is an 82 bp fragment of a *DNAREP1* element. Downstream of this are sections with homology to *Protamine/GD21981*, which include the longest predicted ORF of *Dox*. We designate the 246 bp Dox region with similarity to the 5’ UTR of *Protamine/GD21981* as “2”, and the 474 bp Dox region with coding homology as “3”. Protamines are involved in chromatin compaction during late sperm maturation. In post-meiotic spermatids, histone-based chromatin is replaced by protamines to render a compact, needle-shaped mature sperm (Rathke et al., 2007). Note that the *Dmel-ProtamineA/B* loci (also known as *Mst35Ba/b*) have duplicated locally, whereas *Dsim* and other species in the *melanogaster* subgroup harbor only a single copy at the orthologous region.

Intriguingly, a small portion (63 bp) of the putative Dox ORF is homologous to another *Dmel* gene (*CG15306*), which is not present within *simulans* clade species. We term this segment of Dox as “4” and the longest Dox ORF ends with this unit. Following the internal 359 repeat, the terminal 1080 bp of the *Dox* transcript exhibit homology to *Cubilin* on the X chromosome (termed segment “5”). Thus, the extant *Dox* locus fuses regions of 4 different ancestral protein-coding genes, in addition to various repeat sequences. Next, to understand if there may be other dispersed loci with partial or extended similarities in *Dsim* or *Dmel*, we queried each of these constituent parts of *Dox*. Indeed, this proved to be the case, and we summarize these sequence relationships using percent identity matrices for various Dox sequence features detailed below (**Supplementary Figure 1**).

We located nine matches to segment 1, six of which are located on the X: *Dox*, *MDox*, *GD27797-a* and *GD27797-b*, and two other hits at *CG5004/GD17329*, and *CG15306*. Since the uncharacterized X chromosome loci *GD27797-a/b* share a similar segmental structure as *Dox/MDox*, we name these “*ParaDox*” genes (hereafter, *PDox1* and *PDox2*). The three autosomal matches correspond to one or both arms of the *Nmy* and *Tmy* hpRNAs on 3R (**Figure 1B**). Thus, acquisition of *CG8664/GD15682* sequence appears as an early step during the evolution of Dox family genes.

Segment 2, corresponding to the non-coding portion of the autosomal *Prot/GD21981* gene, hits many of the same loci as segment 1 (**Figure 1C**). We classify these regions of homology to the *Protamine* locus distinctly, as *CG8664/GD15682* and *CG5004/GD17329* contain only noncoding matches to the *Protamine* locus (including portions of the intron), while the four X-linked Dox family loci also match its coding region (segment 3). Protamine homology can also be detected at the Nmy/Tmy hpRNAs. Notably, the four Dox family loci retain clear coding potential that includes the HMG box domain that binds DNA. While not recognized initially (Tao *et al*., 2007a), a similarity search using the Conserved Domain Database (CDDv3.19) identified two significant hits with E-value < 0.001 and includes signature residues of the general HMG box domain (**Figure 1A’**). We can even detect homology between Dox members and human HMG box proteins, emphasizing a likely function for Dox family genes as chromatin factors.

The C-terminal 63 bp of the predicted Dox ORF, termed segment 4, corresponds to sequence from *CG15306* on the X. No orthologous sequence can be found, but homology of *CG15306* to extant Dox family loci suggests likely decimation of the ortholog during Dox family expansion in the simulans clade ancestor. We conclude that insertion of an ancestral Dox family gene at this location disrupted this gene. Sequence similarity with *Dmel CG15306* identified matches at *PDox1*, *PDox2*, *MDox*, and *Dox* on the X, and *Nmy* and *Tmy* hpRNAs on 3R (**Figure 1D**). Finally, segment 5 bears ~1.1kb homology to the C-terminal-encoding region of *Cubilin*. *Cubilin* matches to both *MDox* and *Dox*, but not other Dox family genes or other locations, indicating that this fusion is the most recent integration during the evolution of *MDox/Dox* (**Figure 1E**).

Even with compelling observations of an HMG box domain shared by Dox members, we examined for other possible evidence for translated regions in these genes. As *Cubilin* homologies at *MDox/Dox* are actually located on their antisense strands, any coding potential there would seem to be fortuitous. The *CG8664*-derived segment 1 overlaps the C-terminus of the parental gene, but mostly corresponds to the *CG8664*-3’ UTR. Nevertheless, we find a potential open reading frame (termed ORF13) in this region, and they are aligned in **Supplementary Figure 2**. In addition, copies of a potential open reading frame encoded by protamine-derived segment 2 (ORF5) is aligned in **Supplementary Figure 2**. Although ORF5 is formally from sequence upstream of the protamine transcription unit and/or its 5’UTR, there are more in-frame nucleotide changes and frame-preserving indels across the matching genomic loci, than frame-shifting changes. Neither ORF5 or ORF13 correspond to known domains, and thus their significance is unknown, in contrast to the HMG Box-encoded domains shared by multiple Dox family loci. In any case, the similarity of these putative translated segments provides additional support to the fusion events that generated Dox family genes (**Supplementary Figure 3**).

In summary, *Dox* and *MDox* are members of a larger family of newly-emerged X-linked genes in *Dsim*, which were assembled from pieces of four protein-coding genes that are extant and syntenic in *Dmel*: *CG8664/GD15682*, *Prot/GD21981*, *CG15306*, and *Cubilin*, in addition to 359 satellite repeats (**Figure 1A**). We are particularly intrigued by the fact that multiple X-linked Dox family genes, including two newly-recognized members (*PDox1* and *PDox2*), share an HMG box domain that is derived from protamine. The fact that Dox shares multiple homologies with *Drosophila* protamine hints at a molecular strategy by which it disrupts spermatogenesis, since the replacement of histones by protamines is a key transition in the normal condensation of sperm chromatin during their maturation (Rathke et al., 2014; Wang et al., 2019).

### Tracing the multistep evolutionary origin of *Dox* genes from multiple loci

Given the complex and hybrid structure of *Dox* transcription units, we sought a parsimonious path for their assembly. Our analyses of gene syntenies between *Dmel* and *Dsim* suggest the following model. The key initial event regards how segment 1 from *CG8664/GD15682* might have joined with segments 2 and 3 from *Prot/GD21981* (**Figure 1A**). Intriguingly, all extant similarities to Dox sequence on the X contain adjoining arrangements of segments 1, 2 and 3 (i.e., 1-2-HMG). *Prot/GD21981* are on the 2L arm, while *CG8664/GD15682* are on the X chromosome, and the fusion event likely happened in the *simulans* clade ancestor. Our observations support a model where during the divergence of *simulans* clade from the *Dmel* ancestor, a fusion of these genes from different chromosomes led to the emergence of a chimera. With evidence that protamine gene copies are already in flux (Dorus et al., 2008) (**Figure 2A**), a likely protamine copy mobilized within a *simulans* clade ancestor and inserted within the 3’ UTR of *CG8664*, located on the X chromosome (**Figure 2B**). However, while the contemporary *Dsim* copy of *CG8664/GD15682* contains both segment 1 and segment 2 in its 3’ UTR, it lacks the HMG box-bearing segment 3 (**Figure 2C**). Thus, we infer that the present-day *Dsim* no longer contains the full copy of the original insertion that generated ancestral *Dox* with its motley gang of motifs. We refer to this inferred gene model as the “original gangster *Dox*, *OGDox*” (**Figure 2B**, dotted box). The sublineage of OGDox-related copies that lack the HMG box includes at least one other *Dsim*-specific locus, *GD17329* (the ortholog of *CG5004*) (**Figure 2C, D**). Detailed inspection of these loci indicates complex domain arrangements, which include fragments of the *Tapas* and/or *Krimper* genes (which are both piRNA factors), as well as other transposable elements (**Supplementary Figure 4**).

**Figure 2.**
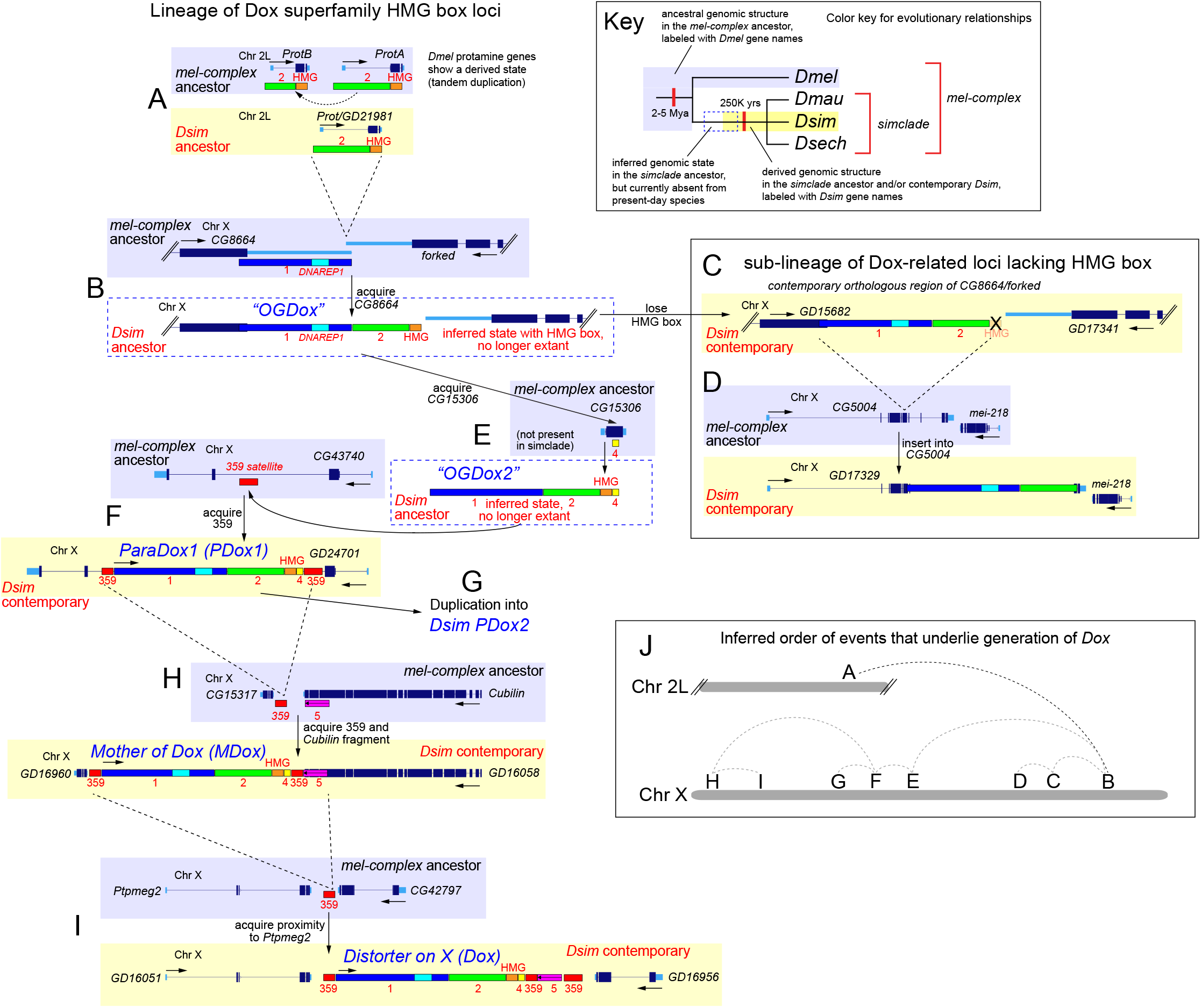
Stepwise origins of *Dox* from a *Protamine*-like ancestor. Upper right, key for the labeling of gene names and structures. Note that many of these regions correspond to extant genomic loci in *Dmel* (purple) and *Dsim* (yellow) but the mobilizations occurred in a *simulans* clade ancestor; they are not meant to indicate that mobilizations occur between contemporary *Dsim* and *Dmel*. In some cases, the inferred events are no longer present in contemporary species (dotted blue boxes). (**A**) In *Dmel*, Protamine shows a tandem duplication (*ProtA/B*), while *Dsim* has a single copy of the Protamine gene (*GD21981*). Inference using an outgroup to *mel*-complex indicated the tandem duplication in *Dmel* is a derived state. Segments from protamine which are acquired in extant Dox superfamily genes are shown in green (segment 2) and orange (HMG) to describe the path to the origin of Dox. (**B**) The first event in the juxtaposition of segments 1-2-HMG is an insertion of Protamine-like gene between *CG8664* (turquoise box in the 3’ UTR of *CG8664* corresponds to piece of DNAREP1 transposable element) and *forked* genes, which we term the “original gangster Dox” (*OGDox*). (**C**) *OGDox* is inferred as an ancestral intermediate, since the contemporary *Dsim* locus lacks the HMG segment and only contains fused segments 1-2. (**D**) Another *Dsim* locus also exhibits segments 1-2 without the HMG box, which derived from an insertion into *CG5004*/*GD17329*. (**E**) The Dox lineage with HMG acquired segment 4 by an insertion of *OGDox* into an ancestral *CG15306* copy. This insertion obliterated *CG15306*, but its relics are identifiable as matches to *Dmel CG15306*. We refer to this insertion into the *Dsim* ancestor as *OGDox2*, again to reflect that it is not retained in present day species. (**F**) Insertion of *OGDox2*, which juxtaposes segments 1-2-HMG-4 into a 359 segment, in the intron of *GD24701* (*CG43740*) yielded the ParaDox (*PDoxA*) gene, which duplicated into two nearly identical dispersed copies in *Dsim* (**G**). (**H**) Mobilization of *PDox* between *Dsim* homologs of *CG15317* and *Cubilin* generated *MDox*. (**I**) *Dox* was generated by mobilization of *MDox* (bearing a fragment *Cubilin*) between *Dsim* ancestors of *Ptpmeg2* and *CG42797*. (**J**) Summary of mobilizations from an ancestral autosomal protamine copy through multiple regions of the X chromosome, which ultimately generate the contemporary *Dox* gene in *Dsim*.

Further evidence of the lability of the inferred *OGDox* locus is the fact that additional copies appear to have mobilized to other genomic locations and splintered further into derivatives that are recognizable by the juxtaposition of segments 1-2-HMG. Their relationships are again clouded by the fact that certain evolutionary intermediates do not appear in present-day genomes. For example, we infer that segment 4 was acquired by insertion of *OGDox* into *CG15306*; however, an ortholog of *CG15306* is no longer extant in the *simulans* clade (**Figure 2E**). Instead, we can now observe that a *de novo* insertion of the gene *GD27797a* bearing segments 1-2-HMG-4 now exists with the intron of *Dsim GD24701* (**Figure 2F**). We note that the ancestral allele, represented by its ortholog *Dmel CG43730*, contains a 359 satellite repeat at the equivalent intronic location. As mentioned, we named this gene “*ParaDox*” and it has duplicated and exists as two nearly-identical copies in *D. simulans* (**Figure 2G**).

*ParaDox* appears to be the parent of *MDox* (**Figure 2H**), which in turn is the parent of *Dox* (**Figure 2I**). We deduce this order, based on the fact that all of these loci share the full complement of 359-1-2-HMG-4-359 segments, but only *MDox* and *Dox* share segment 5, which is related to *Cubilin*. In fact, *MDox* is inserted at *Dsim GD16058*, the ortholog of *Cubilin*, establishing it as the “mother” of *Dox* (**Supplementary Figure 5**). Subsequently, it mobilized between *Ptpmeg2/GD16051* and *CG42797/GD16956* to create *Dox*, which carries a *Cubilin* segment derived from *MDox* and gains a downstream 359 satellite (**Figure 2H, I**). These observations concur with prior observations on *MDox* and *Dox* relationships (Tao *et al*., 2007a).

As we shall see, establishing these complicated mobilization and insertional gymnastics (**Figure 2J**) have implications for interpreting the broader evolutionary dynamics of Dox superfamily genes. First, *ParaDox* appears to have spawned multiple copies, and may represent an ancestral meiotic drive factor. These dynamics are active and different in other species, but their relationships may be traced because gene units containing *Dox* and flanking *Ptpmeg2* are observed in other species. Second, we note that all these Dox superfamily genes are flanked by 359 satellite repeats, and the ancestral syntenic positions of *de novo* insertions also contain 359 blocks (**Figure 2**). This feature may reflect the underlying molecular strategy for amplification of Dox superfamily genes.

### Dox members comprise a rapidly expanding superfamily across three *simulans* clade species

We next analyzed copy number and synteny of Dox superfamily loci from the *simulans* clade sister species *D. mauritiana* (*Dmau*) and *D. sechellia* (*Dsech*). As we noted previously, the repetitive nature of Dox members and their hpRNA suppressors has made them difficult to assemble using short-read or Sanger sequencing (Lin *et al*., 2018); this may be further compounded by the association of many of these loci with satellite repeats (addressed in subsequent analyses). Fortunately, the availability of single-molecule sequencing has recently enabled highly contiguous assemblies of all three *simulans* clade genomes (Chakraborty *et al*., 2021), which we exploited for further analyses.

In *Dsim*, *MDox* is flanked by the *CG15317/GD16960* and *Cubilin*/G*D16058* genes. This *Dsim* arrangement is preserved in *Dmau*, but not *Dsech* (**Figure 3A**). By contrast, while *Dsim Dox* is flanked by *CG42797/GD16956* and *Ptpmeg2/GD16051*, the equivalent genomic regions of *Dmau* and *Dsech* resemble *Dmel*, and lack an intervening *Dox* gene. Thus, the presence of *Dox* in *Dsim* may represent a derived insertion within *Dsim* (**Figure 3A** and **Supplementary Figure 6**). We also observe both conservation and flux for *PDox* genes. *Dsim PDox1* is in the intron of *CG43740/GD24701*, with similar locations of *PDox1* in *Dmau* and *Dsech* (**Figure 3A**). In contrast, *Dsim PDox2* is flanked by *Hk/GD24648* and *CG12643/GD24647*, but comparable regions of *Dmau* and *Dsech* share the ancestral state with *Dmel* (**Figure 3A**). Overall, *Dsim* contains 4 Dox superfamily genes, which we subclassify as Dox family (*Dox* and *MDox*) and ParaDox family (*PDox1* and *PDox2*). Notably, *Dsim PDox* copies have higher homology to Tmy, while *Dox* and *MDox* have higher homology to Nmy (**Supplementary Figure 7**). This suggests that even with our evidence that Tmy has capacity to repress *Dox* and *MDox* (Lin *et al*., 2018), its preferred targets may actually be the PDox genes.

**Figure 3.**
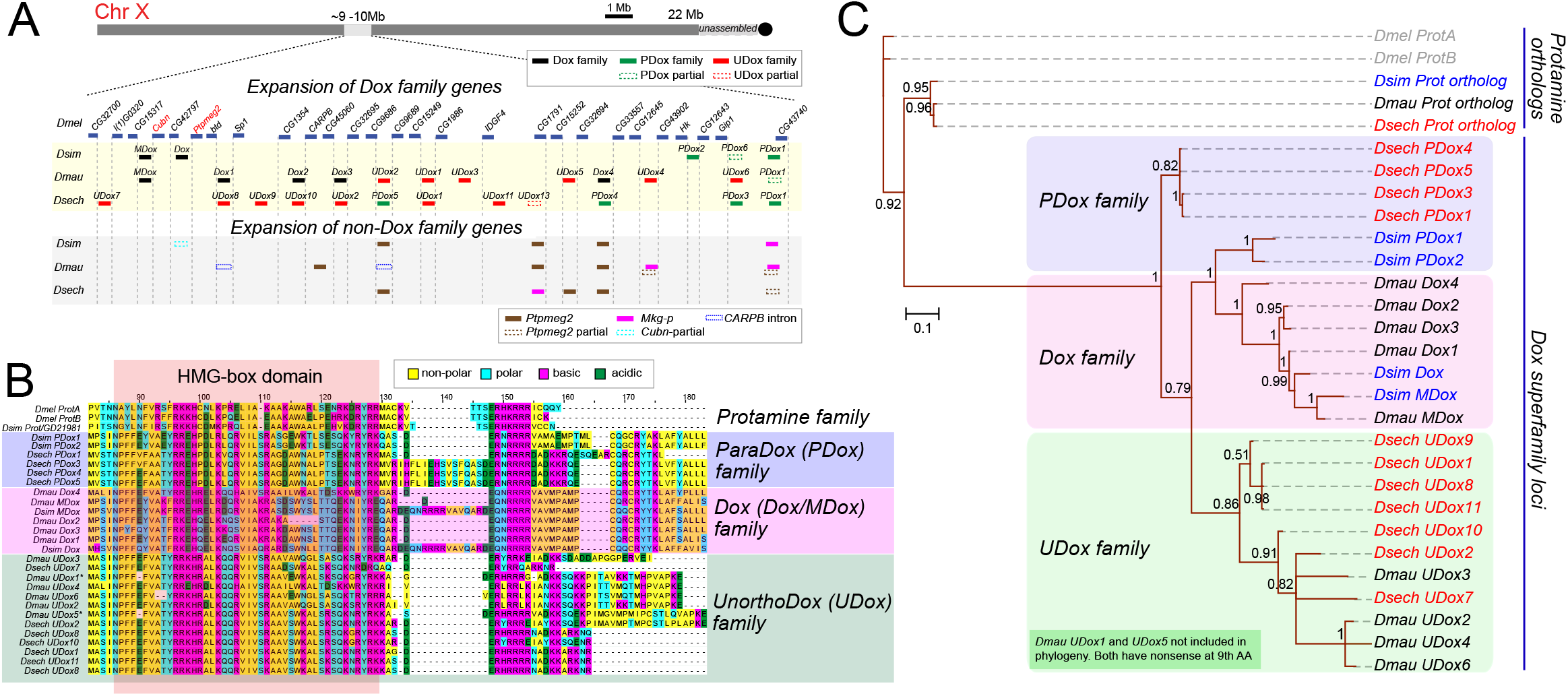
Evolution and diversification of Dox superfamily in the *simulans* clade. (**A**) Chromosomal view of expansion of Dox superfamily and non-Dox family expansions in ~ 1Mb genomic window on the X. Blue tiles show flanking genes as genomic regions to orient expanded copies of Dox superfamily members. (**B**) Classification of Dox superfamily loci into three subfamilies (PDox, Dox, and UDox) was based on amino acid similarity. The highlighted window within the protein alignment indicates the conserved HMG-box domain shared between Protamine and Dox family genes. (**C**) Phylogenetic tree showing similarity relationships among Dox superfamily loci. Highlighted boxes in the tree show clustering of Dox superfamily members into three subfamilies (PDox, Dox, and UDox) based on sequence similarity. Numbers in the tree nodes indicate posterior probability obtained from 1000 bootstrap replicates. *Dmel* ProtA/B and *Dsim* Prot/GD21981 were used as outgroups.

Intriguingly, our homology searches identified massive amplification of Dox superfamily genes in *Dmau* and *Dsech* (**Figure 3A**). We segregated these expansions into different families based on sequence similarity (**Figure 3B**) and their relationship to hpRNAs. In *Dmau*, there are five members of the Dox family, of which only *MDox* is syntenic. These copies have higher homology to hpRNA *Nmy*. In addition, there are six other copies, which have higher homology to an apparent Tmy-like locus (see also later analysis of hpRNA evolution in the *simulans* clade). Because of their distinctive sequences (**Supplementary Figure 9**), which suggest they form a different subfamily, we term these duplicates the UnorthoDox (UDox) family. Although UDox and PDox families share higher homology to Tmy than Nmy, distinct divergence signals cluster these duplications separately (**Figure 3C**). Finally, in *Dsech*, we find four and seven duplicates of the PDox and UDox families, respectively. Details of chromosomal coordinates and sequence features for novel Dox superfamily loci are provided in **Supplementary Table 1**.

Our classification of Dox superfamily genes requires the HMG-box domain, and partial copies lacking HMG-box were excluded from phylogenetic analysis. We find several partial copies lacking HMG, shown as dotted boxes in **Figure 3A** (single partial PDox copies in both *Dsim* and *Dmau*, and one partial UDox copy in *Dsech*). Notably, phylogenetic clustering reveals that most duplicates cluster at the species level, despite synteny at 7 instances. For example, while the syntenic copies of *MDox* from *Dsim* and *Dmau* cluster together, many other syntenic copies show species-level clustering (**Figure 3C**). Two possible scenarios might lead to this pattern. First, independent insertions may have occurred at these sites at species-specific level. Alternatively, gene conversion might occur between copies, resulting in a species-restricted homogenization of copies.

In addition to expansion of Dox family members, we observed expansions of several other gene families associated with Dox superfamily expansion, and have co-amplified in the *simulans* clade as full-length or partial copies. From our testis RNA-seq data, we observe expression of the tyrosine phosphatase *Ptpmeg2* in all three *simulans* clade species, and this gene is syntenic in *Dmel*. In *Dsim*, *Dox* is inserted adjacent to *Ptpmeg2* (**Figure 3A**, **Supplementary Figure 6**). In addition to this syntenic copy there are 3 full length duplicates of *Ptpmeg2* in *Dsim*, and two and three additional full-length copies in *Dmau* and *Dsech*, respectively. Interestingly, similar to Dox superfamily genes, all additional copies of *Ptpmeg2* genes are flanked by 359 satellites and the syntenic, unmodified locations in *Dmel* contain a block of 359 satellite. In addition to full-length copies, we also observe two partial copies of *Ptpmeg2* in *Dmau* and one partial copy in *Dsech* (**Figure 3A**), also flanked by 359 satellites (further discussed in the following section).

Similar to *Ptpmeg2*, we also observe expansion of another gene linked to Dox superfamily expansion. *Mkg-r* is a recently-emerged gene on the X chromosome in the *simulans* clade (Wang et al., 2004), which is a duplicate of the autosomal *Mkg-p* gene. *Mkg-p* encodes a terminal uridyltransferase enzyme related to Tailor, which mediates the uridylation and molecular suppression of splicing-derived pre-miRNAs known as mirtrons (Bortolamiol-Becet et al., 2015; Reimao-Pinto et al., 2015). Notably, while rare mirtrons incorporate into useful regulatory cascades (as evidenced by negative selection), we speculate that they generally comprise a class of rapidly-evolving and potentially selfish regulatory element (Mohammed et al., 2018; Wen et al., 2015b). In addition to a conserved syntenic copy, we find an additional duplicate of *Mkg-r* each in *Dmau* and *Dsech* respectively. Similar to *Ptpmeg2*, the expanded copies of *Mkg-r* are also flanked by 359 satellites (**Figure 3A**). We also note partial matches to *Cubilin* at several locations, and matches to an intronic region of *CARPB* (**Figure 3A**).

Overall, the unusually rapid proliferation and divergence of recently-emerged copies of the Dox superfamily, are atypical for conserved genes. Instead, they conform more closely to the expectation of adaptively evolving genes that often engage conflict scenarios (Agren and Clark, 2018; Zanders and Unckless, 2019). Thus, we speculate that many members of the *simulans*-clade Dox superfamily may be meiotic drivers, and this may also apply to other amplifying genes in this region.

### Dissemination of Dox superfamily genes is tightly linked to insertions into satellite repeats

We were curious as to how the Dox superfamily is capable of such rapid expansion, going from none in *Dmel* to large and highly variable copy numbers in each of the three *simulans* clade species. As mentioned, many Dox superfamily members seem to be flanked by 359 satellites (**Figure 3**). In fact, diverse satellite repeats are known to have highly dynamic copy number in both heterochromatin and euchromatin of *simulans* clade species, and have recently expanded on the X chromosome in the *simulans* clade (Chakraborty *et al*., 2021; Garrigan *et al*., 2014; Khost et al., 2017; Sproul et al., 2020). Moreover, evolution and expansion of satellite sequences have long been suggested to be involved in meiotic drive (Dover, 1982; Walker, 1971).

Amongst the diverse and rapidly evolving sets of *Drosophila* satellite elements, the most abundant and oldest-known class are the 359 element/1.688 satDNA (de Lima et al., 2020; Travaglini et al., 1972). Strikingly, all 29 amplified copies of *Dox* superfamily members in the *simulans* clade are flanked on one or both sides by 359 repeats (**Supplementary Figure 7**). Further inspection reveals distinct modes in the transposition of Dox superfamily genes. Many cases, as exemplified by the inferred movement of *Dsim MDox* to *Dox* (**Figure 2H, I**), involve the localized insertion of a *Dox* gene into a pre-existing 359 satellite, resulting in a flanking arrangement of 359 sequences on either side of the *Dox* gene (**Figure 2I**). For example, we identified examples of such movements that are specific to *Dmau* or to *Dsech*, or that are shared by these species. *MDox* is syntenic and only shared between *Dmau* and *Dsim*, but at the insertion location, a block of 359 satellite repeat is found in both *Dsech* and *Dmel*, indicating insertion sites previously harboring 359 satellite (**Figure 4A**). Similarly, *UDox1* is shared between *Dmau* and *Dsech* and flanked by 359, but in species where *UDox1* is absent, a 359 block is seen at the syntenic location (**Figure 4A**). Details of flanking 359 satellite sequences, and their sequence feature at syntenic locations in *simulans* clade and *Dmel* are provided in **Supplementary Table 2**.

**Figure 4.**
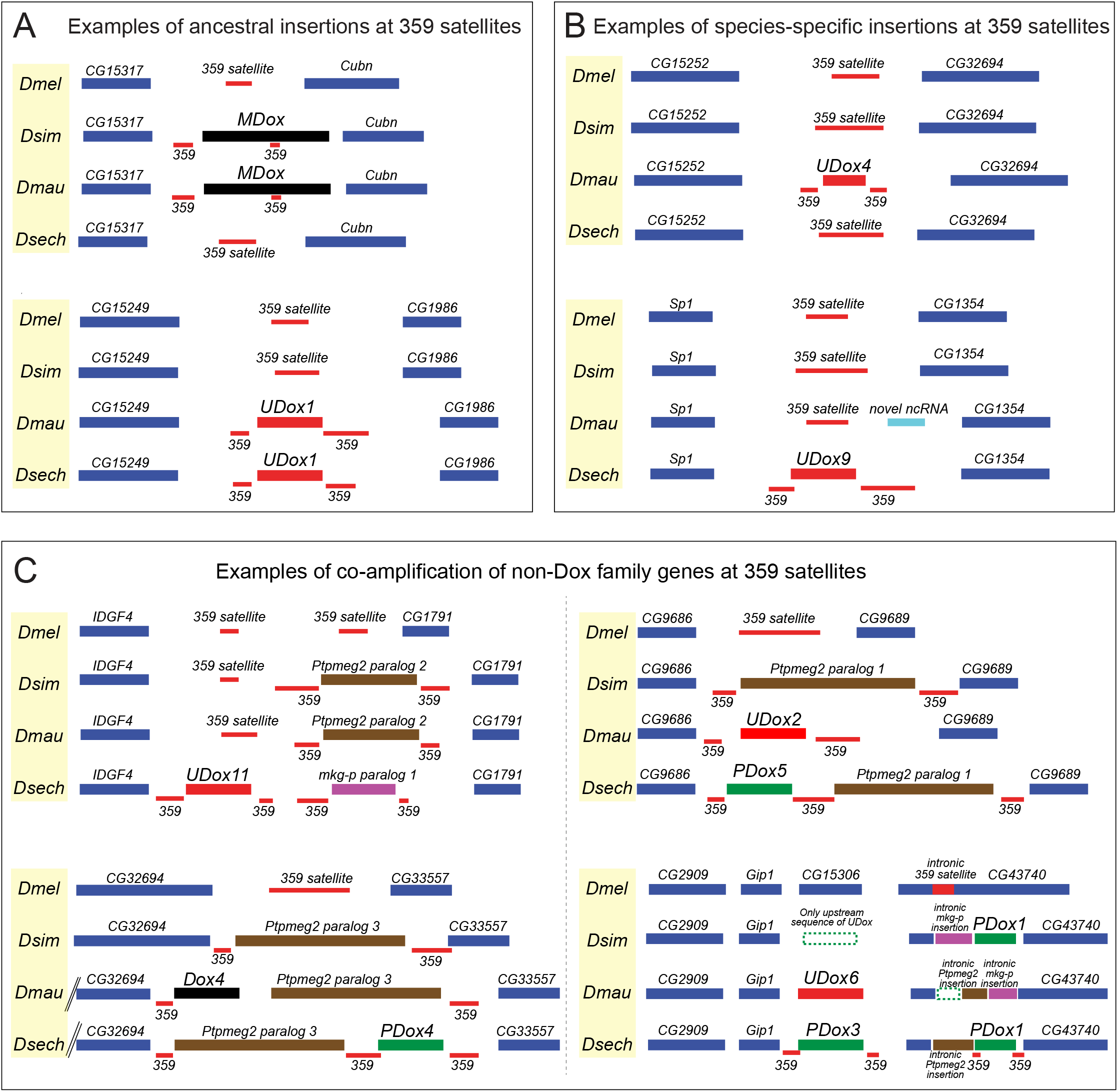
Examples of modes of Dox superfamily expansions in the *simulans* clade. (**A**) Insertion of Dox superfamily loci is associated with 359 satellite repeat. Synteny analysis in the mel-complex revealed 359 sequences at insertion sites are conserved evidenced by their syntenic presence in *Dmel*. (**B**) Within *simulans* clade examples of independent insertions of Dox superfamily members were found indicating their active spread within species. Novel, species-specific insertions/expansions are also associated with 359 satellite repeat at insertion sites, and the synteny of 359 repeat is preserved in other species that lack an insertion. (**C**) Spread of Dox superfamily loci is also linked to co-amplification of two non-Dox family genes on the X chromosome. *Ptpmeg2* and *mkg-p* gene amplification harbors signatures similar to Dox superfamily expansion, where these non-Dox family genes preferentially inserted at 359 satellite regions. *Ptpmeg2* and *mkg-p* co-amplification events show synteny at some instances (insertion between *CG32694* and *CG33557*), but also harbor independent insertions similar to Dox superfamily genes. A detailed synteny analyses of expansions of Dox superfamily with amplification of *Ptpmeg2* and *Mkg-p* are shown in **Supplementary Figure 7**.

The *simulans* clade recently diverged from a shared ancestor ~240,000 years ago, and genome-wide phylogenetic relationships suggested a simple polytomy (Garrigan *et al*., 2012). However, the examples of syntenic insertion found only in two of the three *simulans* clade species, but not all three suggest a more complex scenario at the Dox superfamily region. Of note, inter-specific hybridizations between *simulans* clade species results in hybrid male sterility, but females are fertile and gene flow between these species has been documented (Meiklejohn *et al*., 2018). This can confound species level relationships for these rapidly expanded duplicates. There are also independent insertions into pre-existing 359 satellite blocks in one of these species. For example, *UDox4* is found only in *Dmau*, while *UDox9* appears to be an independent insertion in *Dsech*. All these independent insertion sites also harbor pre-existing 359 satellite, strongly indicating a role for these repeats in mobilization of Dox superfamily genes in the *simulans* clade, putatively via repeat-mediated ectopic recombination (**Figure 4B**).

As mentioned, there is amplification of *Ptpmeg2* and *Mkg-r* copies in *simulans* clade species, often linked to Dox superfamily expansion (**Figure 4C**). The evolutionary dynamics of both of these genes is also tightly associated with 359 repeats, in a manner that can be separated from certain Dox superfamily innovations. For example, *UDox11* insertion between *IDGF4~CG1791* interval which gained a *Ptpmeg2* gene in *Dsim* and *Dmau*, but instead contains a *UDox~mkg-p* cassette in *Dsech*. It is plausible that these may represent independent insertions in the equivalent region of *Dsim/Dmau* vs. *Dsech*, taking advantage of the pre-existing 359 satellite repeats that still exist between *IDGF4~CG1791* in *Dmel*.

Finally, we note seemingly more complex trajectories in which there may have been independent or consecutive insertions into a given genomic locus, given that the three *simulans* clade species can contain all different gene contents between genes syntenic with *Dmel* (**Figure 4C**). However, an alternate scenario is that an entire *Dox~Ptpmeg2* cassette mobilized into this region in a *simulans* clade ancestor, but lost the *Dox* gene itself, followed by subsequent invasion and replacement of this region by a *Dox~mkg-p* cassette (or vice versa). In any case, the tight linkages between 359 satellites and highly variable gene insertions across *simulans* clade species strongly suggest these repeats are causal to this process. Potentially, these repeat sequences might facilitate gene conversion events (Haudry et al., 2020), or perhaps insertions via excised circular DNA copies (Thomas et al., 2014).

### Recurrent emergence of hpRNA suppressors of Dox superfamily loci

With a fuller view of the highly dynamic proliferation of Dox superfamily, we turned to documenting evolutionary strategies for their suppression. We earlier proposed that hpRNA-class siRNA generally derive from their target genes (Wen *et al*., 2015a), and our recent work implies that a major biological implementation of hpRNA regulation is to silence selfish genetic elements (Lin *et al*., 2018). If this is the case, we may predict expansions of Dox superfamily genes to be paired with bouts of corresponding hpRNA births. We previously documented that in *Dsim*, autosomal hpRNAs *Nmy* and *Tmy* are lacking in *Dmel* (Lin *et al*., 2018), and they are not found outside of the *simulans* clade. We now addressed their distribution in *Dsim* sister species *Dsech* and *Dmau*.

In *Dsim*, *Nmy* was proposed to originate via retroposition of *Dox* on chr3R (Tao *et al*., 2007a; Tao *et al*., 2007b), consistent with our general model for hpRNA emergence (Wen *et al*., 2015a). In addition, the frequent association of Dox superfamily genes with repeat sequences provides another mode for the facile generation of additional copies, leading to the birth of multiple hpRNA suppressors to control Dox superfamily genes. In our current analysis, we expand the catalog of hpRNAs that have emerged in concert with Dox superfamily genes, specifically within the *simulans* clade (**Figure 5A**). *Dsim Nmy* is flanked by *GD26005/CG14369* and *GD20491/CG31337*. Synteny analysis showed that *Nmy* is also flanked by these genes in *Dmau*, but no such hpRNA exists in the corresponding location in *Dsech* (**Figure 5B**). The absence of *Dsech Nmy* corresponds to our observation of absence of *Dox/MDox* homologs in *Dsech*. However, the highly abundant PDox copies in *Dsech* (**Figure 3A**) implied that there must be another suppressor of these loci.

**Figure 5.**
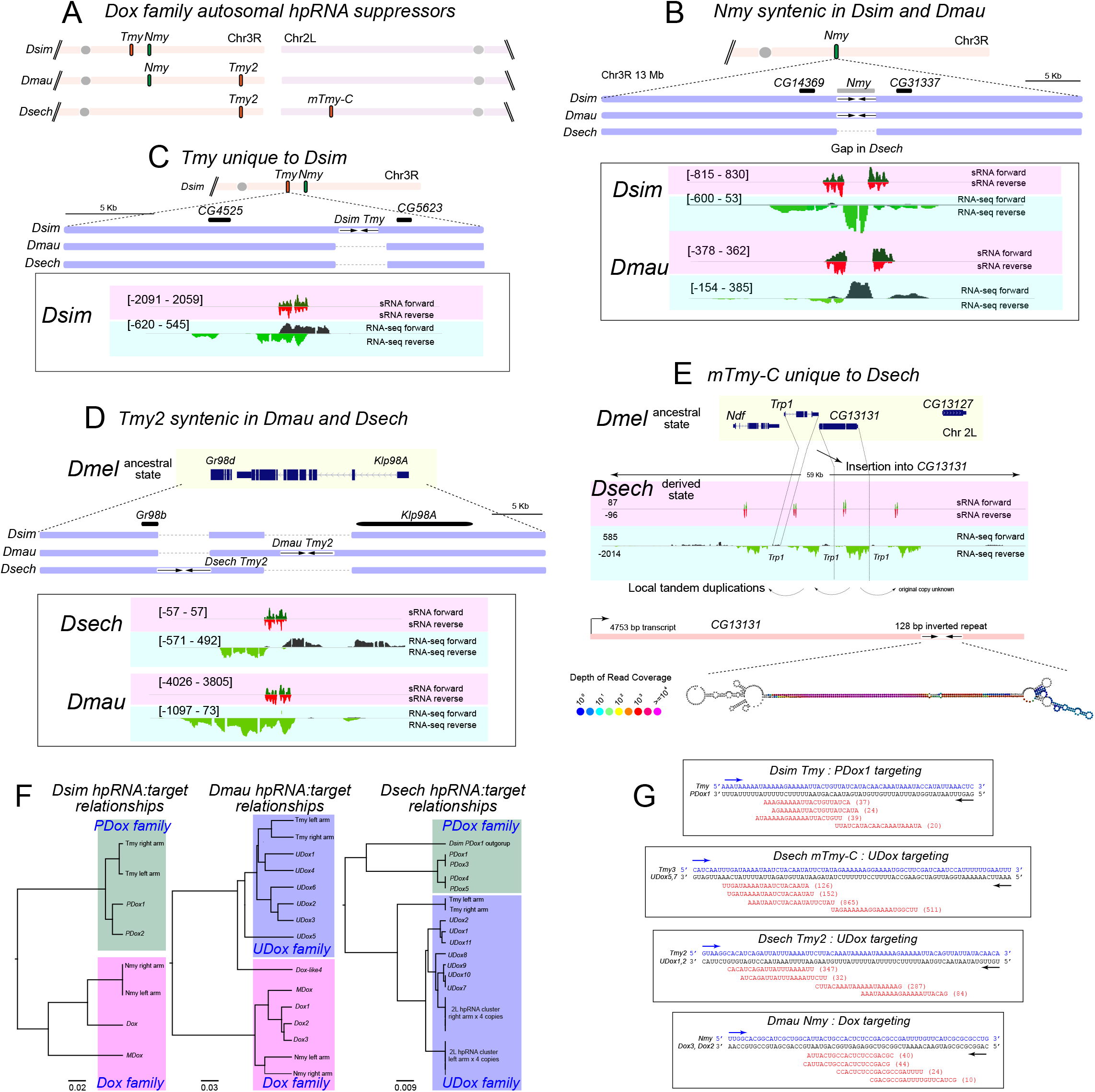
hpRNA:target evolution in the expanded Dox superfamily targeting network. (A) Chromosome map of recently-emerged hpRNAs targeting Dox superfamily in the *simulans* clade. All these hpRNA suppressors are autosomal and reside on Chr3R or Chr2L. (B) Nmy hpRNA is syntenic only in *Dsim* and *Dmau*, but not *Dsech*. Extended nucleotide alignment of flanking sequences in all three species show a gap in *Dsech* indicating emergence of Nmy at this location is syntenic in *Dsim* and *Dmau*. Further evidence for synteny in *Dsim* and *Dmau* was inferred from flanking genes *CG14369* and *CG31337*. (C) Tmy hpRNA is nearby Nmy on Chr 3R and is unique to *Dsim*. Extended nucleotide alignments from all three species at this locus show the presence of *Tmy* only in *Dsim*, and a gap in *Dmau* and *Dsech*. However, the flanking genes *CG4525* and *CG5623* are preserved in all three species. (D) *Tmy2* hpRNA is syntenic between *Dmau* and *Dsech*. *Tmy2* emerged via insertion of a Dox superfamily member between *Gr98d* and *Klp98A*, followed by duplication to generate an hpRNA in the ancestral species. *Dsim Gr89b* and *Klp98A* exhibit *Dmel*-like ancestral state, but these genes are disrupted in *Dmau* and *Dsech* due to hpRNA birth at this locus. (E) Structure and location of novel mini-Tmy-like hpRNA complex (*mTmy-C*) in *Dsech*. Gene models show *Dmel* ancestral state and location of the emergence of Tmy-like hpRNA cluster bearing 4 local duplications. The ancestor to the hpRNA cluster disrupted *CG13131* and in the contemporary state, this hpRNA is flanked by *Ndf* and *CG13127* genes. *CG13131* encodes a 4753 bp transcript and the Tmy-like hpRNA repeat is a 128 bp inverted repeat found at the 3’ end of *CG13131*. Local duplications also affect the flanking gene *Trp1*. RNA-secondary structure for one unit of the Tmy-like hpRNA cluster is shown on the *CG13131* transcript. (F) Phylogenetic trees show that hpRNAs cluster with Dox superfamily targets in the three *simulans* clade species. (G) Example of hpRNA:target relationships, with resulting small RNAs with antisense complementarity to targets. Number of small RNAs observed in *w[XD1]* testis dataset is shown in parentheses. For panels B, C, D, and E, normalized sRNA and RNA-seq tracks depict the structure and expression from hpRNA. Forward strand sRNA reads are shown in dark green and reverse strand sRNAs in red. Similarly, forward strand RNA-seq reads shown in black, and reverse strand RNA-seq reads in light green. Y-axis values indicate normalized read counts for sRNA and RNA-seq.

We next examined *Tmy*, which is located ~2Mb upstream of *Nmy* on Chr 3R in *Dsim*. *Dsim Tmy* is flanked by *GD19044/CG5614* and *GD20331/CG5623*; however, no hpRNA exists in the syntenic region of *Dmau* and *Dsech* (**Figure 5C**). This is consistent with the original introgression genetics whereby the replacement of the *Dsim Tmy* region with its *Dmau* counterpart unleashes meiotic drive phenotypes (Tao *et al*., 2001). Nevertheless, we note that these genetic experiments alone do not fully imply that other species lack *Tmy*, since no molecular analysis was done at the time. It only means that a *Tmy* equivalent does not reside in the syntenic region. The radical amplification of *PDox* and *UDox* paralogs in *Dmau* and *Dsech* prompted us to search more carefully for corresponding hpRNA loci in the contiguous genome assemblies. Indeed, we identified a hpRNA within a syntenic region between *Dmau* and *Dsech*, which generates abundant siRNAs and exhibits extensive homology to Tmy (**Figure 5D**).

Is this *Dmau/Dsech* hpRNA an ortholog, or paralog, of *Dsim Tmy*? We took note of the genes flanking these hpRNAs, and realized they all contain duplicated sequences from a pair of genes, *Gr98d* and *Klp98A*. Interestingly, these genes reside adjacent to each other in the ancestral location shared with *Dmel* (**Figure 5D**). Therefore, we propose that a UDox/hpRNA progenitor inserted between *Klp98A~Gr98d* in a *simulans* clade ancestor, once again taking advantage of flanking satellites. This insertion might have been a tandem copy that was poised to generate small RNAs, but we prefer the scenario in which a three-gene interval containing the UDox progenitor, *Klp98A* and *Gr98d* subsequently duplicated to form a larger genomic inverted repeat. The reason for this is that the *Dmau* and *Dsech* hairpins are not in a precisely syntenic order, but instead reside on the left and right sides of a centrally aligned sequence that is common to *Dsim*. One possibility is that a prior local duplication of the region in a *simulans-clade* ancestor was since collapsed into the single-copy intervals in present-day *Dmau* and *Dsech*, but these resolved via different paths. We call these “*Tmy2*” hpRNAs, and this may have been the source for mobilization from the tip of *Dmau/Dsech* 3R to a more central location of *Dsim* 3R where *Tmy* now resides (**Figure 5A**).

We also took note of the fact that some siRNAs mapped to *Dsech Tmy* also matched to other autosomal locations. This reminded us of our previous discovery of Tmy itself, which we recognized from siRNAs that originally mapped not only to the *Nmy* hpRNA as well as Dox loci on the X, but to an uncharacterized autosomal region that proved to be the Tmy hpRNA (Lin *et al*., 2018). Closer examination of the autosomal cross-mapping reads revealed a repeated locus bearing four tandem copies in *Dsech*. The syntenic region in *Dmel* contains the adjacent *Trp1* and *CG13131* genes. These are still recognizable in *Dsech*, but the *CG13131* copies now contain a ~130bp inverted repeat within its 3’ UTR, which generates siRNAs. The *CG13131~hpRNA~Trp1* multigene unit was subsequently duplicated locally, yielding the present-day disposition in *Dsech* (**Figure 5E**). As these hpRNA inverted repeats are much smaller than Tmy, we refer to this as the *mini-Tmy Complex* (*mTmy-C*).

Clustering of the available hpRNA arms of *Nmy/Tmy/mTmy* loci with Dox superfamily genes in each species reveals a preferred patterns of complementarity (**Figure 5F** and **Supplementary Figures 8 and 9**) which appear to reflect multiple simultaneously active silencing systems that attempt to cope with this bewildering array of putative meiotic drivers. Of note, the newly identified *mTmy-C* loci match well to an diversifying clade of UDox genes in *Dsech*, and are well-positioned to serve as their functional suppressors. Overall, we can find high or indeed perfect antisense complementarity between mature siRNAs from various hpRNAs and individual members of the Dox/PDox/UDox subfamilies (**Figure 5G**).

Since all of these loci are hpRNAs are absent from the closest relative (*Dmel*), they appear to be a prime embodiment of a Red Queen’s race (Van Valen, 1973). Here, the innovation, diversification and amplification of selfish meiotic drive genes of the Dox superfamily, formed by multistep evolution involving the conserved protamine gene, is matched by the innovation of corresponding families of hpRNA-siRNA suppressors. These genetic arms races are segregating in current wild *Drosophila* populations and may be speculated as subject to further ratcheting-up in years to come.

### Rapid evolution of protamine genes within *D. melanogaster*

The analyses presented here may lead to the impression of runaway evolutionary dynamics, unilaterally in the *simulans* clade species compared to *Dmel*. However, since the canonical protamine genes have duplicated in *Dmel* compared to the *simulans* clade, and exhibit signatures of positive selection (Dorus *et al*., 2008), it is clear that protamine loci are subject to recurrent rapid evolutionary dynamics (**Figure 2A**).

We examined the possibility of additional alterations in protamine genes in *Dmel*. Interestingly, queries of the *Dmel* PacBio genome, bearing substantial Y chromosome contigs, revealed multiple copies and pseudogenes of protamine, located within a genomic cluster (**Figure 6A**). This region is adjacent to the 18-member Mst77F cluster located on the Y chromosome (Krsticevic et al., 2015), and was in fact noted as a genomic region (h17 cytoband) containing multiple copies and fragments of several gene families. At the time, these were noted as copies of *CG46192*, *ade5* (*Paics*), *CG12717*, and *Crg-1* (Krsticevic *et al*., 2015). However, subsequent work clarified that CG46192 family and Mst77 family proteins contain the MST-HMG Box domain found in testis-restricted proteins (Doyen et al., 2015). Of note, both Mst77F and protamines replace histones during compaction of sperm chromatin (Jayaramaiah Raja and Renkawitz-Pohl, 2005). We find that CG46192, along with its cluster copies and pseudogenes, are more similar to protamine than Mst77F/Mst77Y proteins (**Figure 6B**), indicating that they represent a distinct amplification event. Moreover, there is a complex history to the emergence of this cluster, since *Paics* and *CG12717* are adjacent genes on the X chromosome while protamine (*Mst35Ba/b)* genes are located on chr 2L (https://flybase.org/). Although perhaps coincidental, we note that *CG12717* encodes a phosphatase, while *simulans* clade species have *de novo* amplifications of a different phosphatase (*Ptpmeg2*); *Drosophila* species contain an unusually large number of phosphatases, which include many rapidly evolving copies (Miskei et al., 2011). The assembled *Dsim* Y does not appear to contain copies of MST-HMG Box genes.

**Figure 6.**
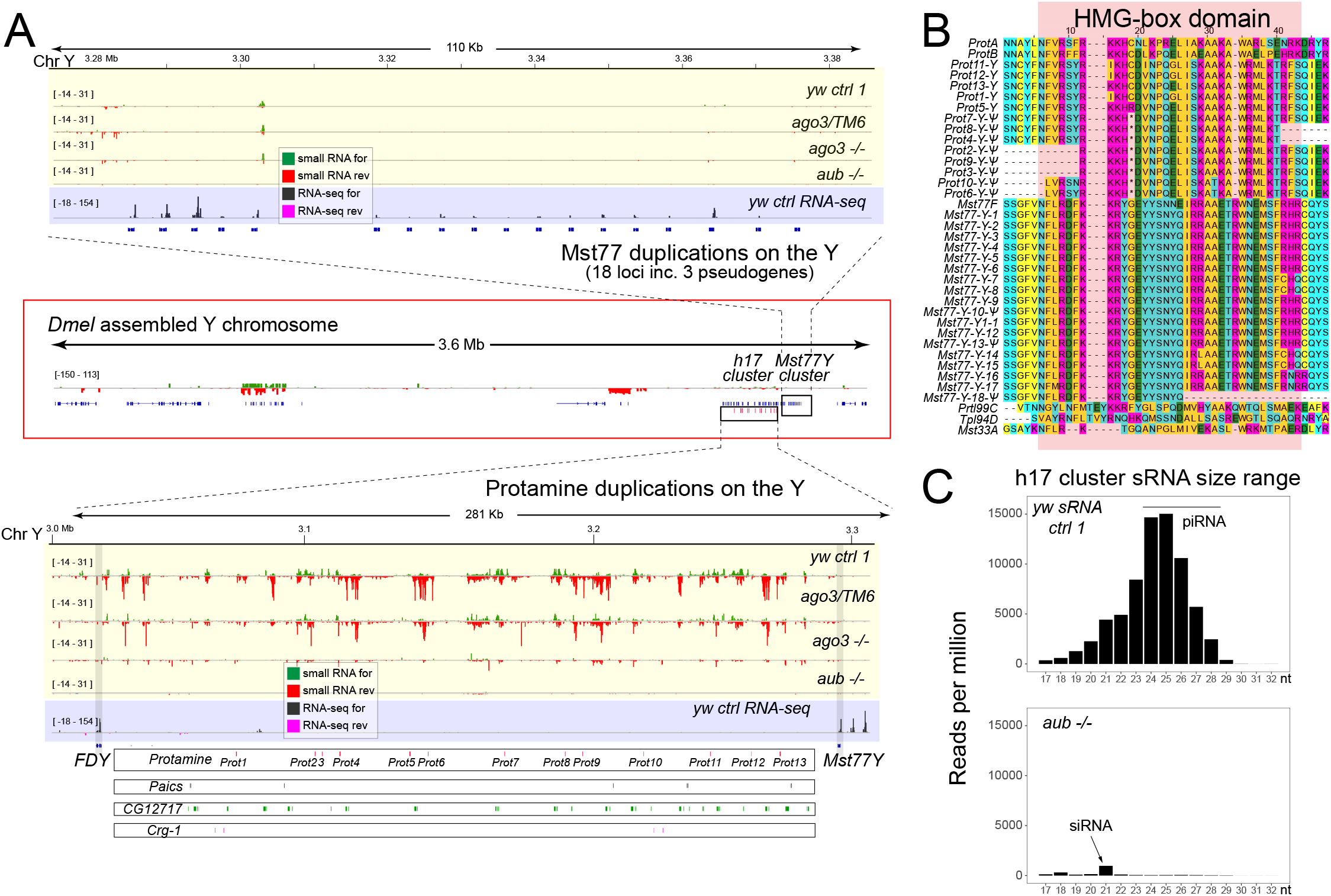
Genomic expansion of protamine genes in *D. melanogaster*. As noted, the autosomal protamine locus is in a derived state in *Dmel*, as it is locally duplicated, unlike *simulans* clade and outgroup *Drosophila* species (**Figure 2A**). (A) Genome browser tracks of small RNA data (yellow) and RNA-seq data (purple) from control (ctrl) and ago3 heterozygous (over TM6) testis, as well as from piRNA pathway mutant testis (*ago3* and *aubergine/aub*). Two regions of the assembled Y chromosome (central red box) are shown as enlargements. Top, expansion of Mst77 genes. Protamine belongs to the MST-HMG box family, for which the autosomal member Mst77F was previously observed to have broadly expanded on the Y chromosome (Mst77Y cluster). Bottom, adjacent to the Mst77Y cluster, in the h17 cytoband, is another cluster bearing repeated portions of multiple protein-coding genes, including protamine. The annotated genes in the Mst77Y cluster are associated with testis RNA-seq evidence, but not small RNA data. By contrast, the h17 cluster is associated with abundant small RNA data, but not RNA-seq data. Most of these reads seem to be piRNAs, since they are depleted in piRNA mutant testes. (B) Alignment of MST-HMG box members indicates that the h17 copies on the Y are more closely related to Protamine than to Mst77 members or other MST-HMG box members. (C) Small RNAs from the h17 cluster are mostly piRNA-sized (~23-28 nt in these data) and are depleted in testis mutated for the piRNA factor *aubergine* (*aub*). The minor population of h17 cluster small RNAs in *aub* mutant testis are siRNA-sized (21 nt).

To address if there may be relationship of the h17 cluster with small RNAs, we examined wildtype testis data with that of the piRNA factor *aubergine* (*aub*) (Nagao et al., 2010). Interestingly, abundant small RNAs map to the h17 Y cluster (**Figure 6B**), but not to the adjacent Mst77Y cluster (**Figure 6A**). However, these are dominantly in the piRNA-sized range (**Figure 6C**). Evidence that these are in fact piRNAs comes from the fact that their accumulation is strongly decreased in *aubergine* mutant testis (**Figure 6C**). The observation of abundant testis piRNAs from the h17 cluster was independently reported while this work was in revision (Chen et al., 2021). Interestingly, the small amount of remaining h17 cluster small RNAs in these mutants are 21nt in size (**Figure 6C**), suggesting an interplay of piRNA and siRNA biogenesis at this cluster, as seen for other piRNA clusters in *D. melanogaster* (Lau et al., 2009; Malone et al., 2009).

## Discussion

### Rapid evolutionary dynamics of *de novo* Dox family meiotic drive genes and origin from protamine

In this study, we reconstruct the ancestry and diversification of an expanded family of Dox genes and their presumed hpRNA/siRNA suppressor loci. These genes exhibit partly overlapping content amongst the three *simulans* clade species, but exhibit numerous unique genomic copies and innovations within each species. Notably, all of these Dox family loci are absent from their closest sister species *D. melanogaster* and other species in the *Dmel* group. This implies the birth of a meiotic conflict in the *simulans* clade ancestor, and subsequent cycles of proliferation of Dox family drivers and their subsequent suppression by hpRNA loci (summarized in **Figure 7**).

**Figure 7.**
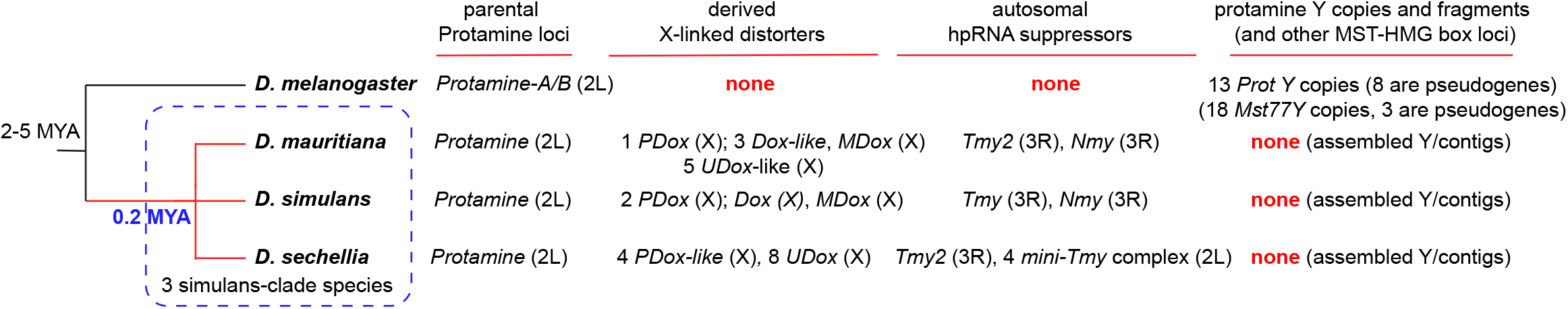
Summary of expansion of the Dox superfamily in *simulans* clade fruitflies. A) Phylogenetic relationship between *Dmel* and the *simulans* clade. The *simulans* clade diverged from *Dmel*-like ancestor ~2-5 MYA, while individual *simulans* clade species diverged ~0.2 MYA. The parental gene for all Dox superfamily copies and their hpRNA suppressors is Protamine in 2L. *Dmel* Protamine exhibits an additional local duplicate, along with multiple *de novo* copies on the Y; however, this species lacks any Dox superfamily loci or corresponding hpRNAs. Amongst the three simulans clade species, there are no apparent Y-copies of protamine, but instead highly variable numbers and classes of Dox superfamily genes and homologous hpRNAs.

Until now, the presence of any distinctive nucleotide content of *Dox* was unknown, other than its homology to *MDox* and the hpRNA loci *Nmy* and *Tmy* (Lin *et al*., 2018; Tao *et al*., 2007a; Tao *et al*., 2007b). However, the recognition of multiple potential open reading frames that are shared with other genomic sources, and their syntenies amongst *simulans* clade species and *D. melanogaster*, allowed us to trace stepwise origins of an ancestral *Dox* gene from multiple genomic regions that remain identifiable in *D. melanogaster*. Importantly, this includes insertion of an ancestral protamine copy into another locus to yield a hybrid gene product, which subsequently exploited flanking satellite context to undergo amplification and mobilization. The functions of these genes have yet to be directly analyzed beyond the finding that *dox* mutations suppress the female-biased progeny sired by *nmy* hpRNA mutant males (Tao *et al*., 2007a). Nevertheless, the rapid diversification of Dox family genes, which assort into at least three recognizable subfamilies (and potentially more, depending on the granularity of subdivision), suggests that many members of this family participate in meiotic drive.

The notion that multiple Dox genes encode selfish activities is further supported by the coevolution of multiple hpRNA loci that exhibit preferential complementarity to Dox subfamilies. Of note, we previously documented that hpRNAs are always evolutionarily young, but evolve adaptively with their targets (Wen *et al*., 2015a). Thus, they are ideal weapons to silence incipient selfish loci, a notion well-supported by their evolutionary and sequence characteristics. It is known that mutants of the individual, *de novo* hpRNA Nmy exhibit highly deleterious phenotypes in spermatogenesis (Lin *et al*., 2018; Tao *et al*., 2007b), which contrasts with the generally subtle defects of scores of highly conserved miRNA genes in *Drosophila* (Chen et al., 2014). Although genetic analyses of other *Nmy*, *Tmy* family and *mini-Tmy* hpRNAs awaits, we predict they will also affect the development of the male germline in some fashion.

It is presently unclear whether Dox family genes might encode multiple proteins, as suggested by the seemingly operon-like organization of most family members with possible additional ORFs (**Supplementary Figure 2**). We note that “ORF5” can be recognized in the ancestral protamine gene of *D. melanogaster* but its contribution to the phenotype of targeted deletions of the adjacent *ProtamineA/B* (Mst35Ba/b) genes has not been tested (Tirmarche et al., 2014). It is possible that ORF13 and ORF5 do not serve a trans-acting function, but are simply evolutionary relics of the stepwise acquisition of genomic pieces during the emergence of an ancestral *Dox* gene. Another possibility is that they serve a regulatory role in cis as upstream ORFs, to modulate the output of the HMG Box ORF. As of now, we are quite intrigued with this clear domain homology, since dozens of Dox superfamily genes are homologous to the HMG-Box domain found in protamines, with residues shared in even more distant relatives that encode transcription factors (Sox genes).

Sperm chromatin becomes highly condensed during maturation, coinciding with the replacement of histones with protamines, both in flies (Rathke *et al*., 2014) and mammals (Wang *et al*., 2019). Since sex chromosome conflict is most apparent in the male germline, the homology of Dox family proteins to protamine provides a testable foundation for understanding their role in meiotic drive systems that distort fidelity and quality of spermatogenesis, namely Winters and Durham drive (Tao *et al*., 2001; Tao *et al*., 2007b). Interestingly, the most well-known meiotic drive system in flies, the autosomal *Segregation Distorter (SD)* system of *D. melanogaster* (Larracuente and Presgraves, 2012), was recently linked to protamine dysfunction (Herbette et al., 2021), and protamine knockdown also enhances SD drive capacity (Gingell and McLean, 2020). We note that while the histone-to-protamine transition can occur in *D. simulans nmy* mutants (Yasuno et al., 2013), the stronger condition of *D. simulans* RNAi mutants (which compromise all hpRNAs) strictly block this process since histones remain associated with *dcr-2* and *ago2* mutant spermatids (Lin *et al*., 2018). Therefore, we hypothesize that the histone-to-protamine transition is a sensitive point in spermatogenesis that may be under recurrent attack by meiotic drivers and selfish genetic elements. Indeed, the intimate connection of protamines and sex chromosome conflict is bolstered by the independent expansion of euchromatic and Y chromosome protamine copies in *D. melanogaster*.

### Evolutionary lability of selfish element and their suppressor systems

It was recently documented that, despite an overall low amount of gene flow between *D. mauritiana* and *D. simulans*, including on the X, the *Dox/MDox* interval recently transferred between these species (Meiklejohn *et al*., 2018). Thus, this represents one mechanism for the spread of selfish genetic elements between related species. However, we are struck by the highly dynamic proliferation and diversity of Dox family loci amongst the three quite closely related *simulans* clade species, which indicates that gene flow cannot account for their evolution. Our observation of near-universal existence of 359 satellite sequences flanking most Dox superfamily genes strongly suggests that these are involved in their evolutionary strategy of dispersal. This notion is further bolstered by the existence of satellite-flanked multigene units bearing a Dox family gene, and even their existence surrounding hpRNA genes.

Nonallelic homologous exchanges involving repeats are a source of genomic instability, including deletion, amplification and translocation. Therefore, it may be advantageous for a selfish gene to be embedded in the context of repeats. However, since the landscape of Dox genes includes substantial numbers of dispersed copies, these are not necessarily explained by changes in local tandem copy numbers. Gene conversion often takes place locally, although it is known to occur in an interlocus fashion (Chen et al., 2007). However, we note that satellite repeats not only exhibit highly distinctive profiles between species, and even amongst individuals, they are known to be associated with the formation of extrachromosomal circular DNA (eccDNA) via intrachromatid recombination (Mourier, 2016). The existence of satellite-associated eccDNA has been observed in diverse species, ranging from plants (Navratilova et al., 2008), flies (Cohen et al., 2003), and mammals (Cohen et al., 2006). Recent surveys and analyses of *simulans* clade species further confirm high rates of genomic turnover and innovation of satellite sequences and their abundant representation in eccDNA (de Lima *et al*., 2020; Sproul *et al*., 2020; Wei et al., 2018).

The 359 satellite (also known as 1.688 satDNA) is the evolutionarily oldest and also the most abundant *Drosophila* satellite sequence (de Lima *et al*., 2020; Travaglini *et al*., 1972). Precise analyses of the genomic makeup of repeat sequences, including satellites, are generally difficult due to their misassembly in short-read sequenced genomes; yet it was recognized some time ago that 359 satellites have recently expanded on the X chromosomes of *simulans* clade species (Garrigan *et al*., 2014). The advent of single-molecule long read sequencing has enabled much greater precision in documenting the high rate of evolutionary dynamics of 359 and other satellite sequences across *simulans* clade species (Chakraborty *et al*., 2021; Khost *et al*., 2017; Sproul *et al*., 2020). Thus, Dox superfamily loci may potentially hijack the intrinsically elevated evolutionary dynamics of satellite sequence to fuel their spread and amplification, potentially involving exchanges to and from the extrachromosomal pool.

These principles may apply more generally. Similar to the rapid amplification of Dox superfamily genes, and their association with 359 satellite repeats, recent findings from meiotic drive families across a range of non-*Drosophila* species recapitulate two broad themes: (1) ampliconic nature of drive loci, for example *Slx/Sly* in mice (Cocquet et al., 2012; Cocquet et al., 2010; Moretti et al., 2020), *Spok* meiotic drive gene family in *Podospora anserina* (Vogan et al., 2019), and the *wtf* meiotic drive gene family in *S. pombe* and certain other yeasts (Hu et al., 2017; Nuckolls et al., 2017), and (2) association of meiotic drive gene family expansion with repetitive elements, for instance, the expansions of Spok family genes mediated by Enterprise-class transposable elements (Vogan et al., 2021).

Finally, we document hpRNA/siRNA suppressors of meiotic drive loci, and we previously documented that hpRNAs evolve very rapidly in general, and seem to exist only for short evolutionary periods. We also expand the scope of protamine-class innovations and expansions between closely-related *simulans*-clade species, and potentially linking independent evolution of protamine genes in *Dmel* to the formation of a *de novo* piRNA cluster. While this work was in review, Aravin and colleagues reported thematically related findings and suggest similar evolutionary lability of target control by siRNA and piRNA pathways, even between closely related species (Chen *et al*., 2021). Their study details that the *Dmel* h17 region is a Y chromosome piRNA cluster expresed in testis, which exerts particularly strong repression of the ancestral *CG12717* copy, which encodes a SUMO protease. However, in the *simulans*-clade species *Dmau*, *CG12717* is instead under siRNA control (Chen *et al*., 2021). These findings further emphasize the notion that particular collections of genes evolve with extraordinarily rapid and recurrent dynamics in the testis, and that some of these are targeted by small RNAs, implying scenarios of conflict regulation.

## Materials and Methods

### Genome data

PacBio genome data for *simulans* clade species (Chakraborty *et al*., 2021) was obtained from SRA through the Bioproject ID: PRJNA383250. Individual genome assemblies for *D. simulans*, *D. mauritiana*, and *D. sechellia* are available through genome assembly IDs: ASM438218v1, ASM438214v1, and ASM438219v1 respectively.

### RNA-seq library preparation and sequencing

We used *Dsim w[XD1]* and *Dmau w[1]* 14021-0241.60, which were used for PacBio genome sequencing (Chakraborty *et al*., 2021), and Dsech 14021-0248.25 used for the short read genome (Clark et al., 2007). We isolated total RNA from ~ 5-day-old flies and for *Dsim, Dmau and Dsech* samples. We extracted RNA from testes (dissected free of accessory glands) using Trizol (Invitrogen). We made two independent dissections to generate biologically replicate RNA samples, whose quality was assessed by Bioanalyzer. We used the Illumina Truseq Total RNA library Prep Kit LT to make RNAseq libraries from 650 ng of total RNA. Manufacturer’s protocol was followed except for using 8 cycles of PCR to amplify the final library instead of the recommended 15 cycles, to minimize artifacts caused by PCR amplification. All samples were pooled together, using the barcoded adapters provided by the manufacturer, over 2 flow cells of a HiSeq2500 and sequenced using PE75 at the New York Genome Center.

### sRNA library preparation and sequencing

For small RNA analysis, we extracted RNA from testes and accessory glands of 7-day-old *Dsim w[XD1]*, *Dmau w[1]* 14021-0241.60, and *Dsech* 14021-0248.25 strains using Trizol (Invitrogen). 1 μg of total RNA were used to prepare small RNA libraries as described (Lee and Yi, 2014), with the addition of QIAseq miRNA Library QC Spike-ins for normalization (Qiagen). Adenylation of 3’ linker was performed in a 40 μL reaction at 65°C for 1 hr containing 200 pmol 3’ linker, 1X 5’ DNA adenylation reaction buffer, 100 nM ATP and 200 pmol Mth RNA ligase and the reaction is terminated by heated to 85°C for 5 min. Adenylated 3’ linker was then precipitated using ethanol and was used for 3’ ligation reaction containing 10% PEG8000, 1X RNA ligase buffer, 20 μM adenylated 3’ linker and 100 U T4 RNA Ligase 2 truncated K227Q. The 3’ ligation reaction was performed at 4°C overnight and the products were purified using 15% Urea-PAGE gel. The small RNA-3’ linker hybrid was then subjected to 5’ ligation reaction at 37°C for 4 hr containing 20% PEG8000, 1X RNA ligase buffer, 1 mM ATP, 10 μM RNA oligo, 20 U RNaseOUT and 5 U T4 RNA ligase 1. cDNA synthesis reaction was then proceeded immediately by adding following components to the ligated product: 2 μl 5x RT buffer, 0.75 μl 100 mM DTT, 1 μl 1 μM Illumina RT Primer, and 0.5 μl 10 mM dNTPs. The RT mix was incubated at 65 C for 5 min and cooled to room temperature and transfer on to ice. 0.5 μL of superscript III RT enzyme and 0.5 μL RNase OUT were added to the RT mix and the reaction was carried out at 50°C for 1 hr. cDNA libraries were amplified using 15 cycles of PCR with forward and illumine index reverse primers and the amplified libraries were purified by 8% non-denaturing acrylamide gel. Purified libraries were sequenced on HiSeq2500 using SR50 at the New York Genome Center. Oligo sequences and reagents used for library preparation are provided in Table A1.

### Gene annotations and *de novo* transcriptome annotation

We used the UCSC liftover coordinate conversion tool to obtain PacBio gene annotations using *D. simulans* r2.02 Flybase annotations. Briefly, lastz tool from UCSC was used to construct pairwise genome alignments between r2.02 and PacBio genome assemblies. Next axtChain tool implemented in KentUtils (https://github.com/ucscGenomeBrowser/kent) was used to generate a chains file for individual chromosomes to compare alignments and coordinates between two genome assemblies. Then, using the liftOver tool in KentUtils we converted gene annotations in gtf/bed format from r2.02 to an equivalent PacBio gtf/bed file for browsing on IGV/UCSC genome browser. We performed *de novo* transcriptome annotation from testis RNA-seq data in *Dsim*, *Dmau* and *Dsech* using Cufflinks (Trapnell et al. 2010, 2012). *De novo* transcriptome annotation was then overlapped with known gene annotations from liftOver files using intersectBed implemented in bedtools (https://bedtools.readthedocs.io). Non-overlapping annotations were then added to the gtf file as novel testis transcripts in each species. Most *Dmel* analyses used Flybase dm6, but for *Dmel* Y chromosome analysis, we used an improved Y assembly that incorporates additional contigs absent from dm6 (Chang et al., 2019).

### Repeat annotation

PacBio repeat annotation of *Dsim* clade genomes was performed as described (Chakraborty *et al*., 2021). Briefly, new complex satellites were annotated using Tandem repeat finder; and novel TEs were annotated using the REPET TE annotation package (Flutre et al. 2011). The resulting novel satellite and TE sequences were added to the curated *Dmel* Repbase library and repeats in the PacBio genome assemblies were annotated using RepeatMasker v4.0.5. Annotated repeat tracks were then split into TEs, and simple repeats and converted into BED format for viewing on IGV.

### Data analysis

RNA-seq data: Paired-end RNA-seq reads was mapped to both PacBio genome assemblies for *Dsim*, *Dmau*, and *Dsech* using hisat2 aligner with the following command hisat2-x indexed_genome_assembly -1 $ read1.fastq.gz -2 read2.fastq.gz -S file.sam. The alignment file in SAM format was then converted to a compressed BAM file using SAMTOOLS (Li et al. 2009) with the following commands: 1) samtools view -bS file.sam > file.bam 2) samtools sort file.bam > file_sorted.bam 3) samtools index file_sorted.bam. Mapping statistics for the BAM alignment files were obtained using bam_stat.py script from the RSeqQC package (Wang et al., 2012).

small RNA data: sRNA reads were processed as follows: Raw sequence reads were adapter trimmed using Cutadapt software (https://cutadapt.readthedocs.io/en/stable/). After clipping the adapter sequence, we removed the 4bp random-linker sequence inserted at 5’ and 3’ of the sRNA sequence (total 8bp). After filtering <=15 nt reads, we mapped the small RNA data to PacBio genome assemblies using Bowtie (with –v0 –best –strata options.) The resulting small RNA alignments in SAM format were converted to BED for downstream processing using the BEDops software and visualized on IGV.

### Phylogenetic analysis

For phylogenetic analysis of Dox superfamily genes, we constructed an alignment of CDS for each ortholog using the translation align feature in Geneious (version 11.0.4). For this alignment, we excluded two UDox copies (*UDox1* and *UDox5*) in *Dmau*, which appear to carry premature stop codons. The alignment was performed using the Geneious multiple alignment sequence feature using the global alignment with free end gaps. For the alignment, a 65% similarity (5.0/-4.0) cost matrix was used with the following gap penalty parameters: (1) Gap open penalty: 12, (2) Gap extension penalty: 3, and (3) Refinement iterations: 2. The resulted alignment was then manually curated to ensure proper alignment. Phylogenetic analysis on the nucleotide alignment was performed using MrBAYES (Huelsenbeck and Ronquist, 2001) plugin in Geneious software v11.0.4. For this analysis, we used the HKY85 substitution model with *Dmel* ProtA/B as an outgroup. A gamma rate variation option was used with the following gamma categories: 4. For Monte Carlo Monte Chain (MCMC) settings, we used the following parameters: (1) chain length: 10000, (2) Heated chains: 4, (3) Heated chain Temp: 0.2, (5) Subsampling Frequency: 100, (6) Burn-in length: 1000, (7) Random seed: 7826. For priors, we used the Unconstrained Branch Lengths option with the following GammaDir parameters (1,0.1,1) and the Shape Parameter: Exponential (10).

### Homology and domain searches

Sequence homology search for putative ORFs encoded in the Dox transcript and search for Dox like sequences in the PacBio assemblies were performed using command-line version of blastn and/or tblastn implemented in BLAST 2.2.31+ (Altschul et al., 1990). Search for conserved protein domains in the Dox family genes was performed using both HMMER v3.3.2 and NCBI conserved domain database (CDD v3.19). In addition to these search algorithms, we also used the Phobius domain search tool (https://phobius.sbc.su.se/), which specifically searches for similarities to signal peptides and transmembrane topology.

## Data availability

Paired-end RNA-seq reads from *Dmau*, *Dsim*, *Dsech* testis, and small RNA data from *Dmau* and *Dsech* are under submission to GEO.

## Acknowledgments

We thank Jiayu Wen for initial investigations into the domain structure of Dox, and Peter Smibert for the first recognition that Dox/MDox bear homology to protamine. Work in the ECL group was supported by the National Institute of General Medical Sciences (R01-GM083300) and National Institutes of Health MSK Core Grant P30-CA008748, and US-Israel Binational Science Foundation BSF-2015398.

